# Targeting Toxic Nuclear RNA Foci with CRISPR-Cas13 to Treat Myotonic Dystrophy

**DOI:** 10.1101/716514

**Authors:** Pornthida Poosala, Sean R. Lindley, Kelly M. Anderson, Douglas M. Anderson

## Abstract

Human monogenetic diseases can arise from the aberrant expansion of tandem nucleotide repeat sequences, which when transcribed into RNA, can misfold and aggregate into toxic nuclear foci^1^. Nuclear retention of repeat-containing RNAs can disrupt their normal expression and induce widespread splicing defects by sequestering essential RNA binding proteins. Among the most prevalent of these disorders is myotonic dystrophy type 1 (DM1), a disease occurring from the expression of a noncoding CTG repeat expansion in the 3’UTR of the human *dystrophia myotonica protein kinase* (DMPK) gene^2,3^. Here we show that RNA-binding CRISPR-Cas13, with a robust non-classical nuclear localization signal, can be efficiently targeted to toxic nuclear RNA foci for either visualization or cleavage, tools we named *hilightR* and *eraseR*, respectively. *HilightR* combines catalytically dead Cas13b (dCas13b) with a fluorescent protein to directly visualize CUG repeat RNA foci in the nucleus of live cells, allowing for quantification of foci number and observation of foci dynamics. *EraseR* utilizes the intrinsic endoribonuclease activity of Cas13b, targeted to nuclear CUG repeat RNA, to disrupt nuclear foci. These studies demonstrate the potential for targeting toxic nuclear RNA foci directly with CRISPR-Cas13 for either the identification or treatment of DM1. The efficient and sequence programmable nature of CRISPR-Cas13 systems will allow for rapid targeting and manipulation of other human nuclear RNA disorders, without the associated risks of genome editing.

---

Myotonic dystrophy type 1 (DM1) is an autosomal dominant human monogenic disease characterized by progressive myotonia, muscle wasting, cardiac arrhythmias, and cognitive dysfunction^4^. DM1 is the most common form of adult-onset muscular dystrophy, occurring in roughly 1 in 8,000 individuals^5^. Mutant DMPK mRNAs with greater than ∼50 CUG repeats form toxic nuclear RNA foci, which prevent normal DMPK expression and induce widespread defects in alternative splicing and polyadenylation by sequestering members of the muscleblind-like (MBNL) family of RNA binding proteins^6-13^. Loss of MBNL1 and 2 function account for up to 80% of the observed DM1 phenotype, however, overexpression of MBNL proteins have had limited success rescuing splicing defects and only modest amelioration of muscle function in DM1 mouse models, suggesting a multifactorial cause for DM1 symptoms^14-17^. The multitude of dysfunctional pathologies associated with the formation of toxic RNA foci suggests that targeted degradation of CUG expansion (CUG^exp^) RNA would provide therapeutic benefit to DM1 patients.

Despite intense effort, there are no currently approved therapies designed to treat DM1. Antisense oligonucleotides (ASOs) targeting CUG-repeats have been successfully used to reduce the levels of toxic RNAs and disrupt binding and sequestration of MBNL proteins in animal models^18,19^. However, significant challenges remain for the delivery of these molecules at therapeutically effective levels in human skeletal muscle. Genome editing approaches using CRISPR-Cas9 DNA endonucleases have been employed to directly edit the DMPK gene locus. However, CRISPR-Cas9 editing at either the 5’ or 3’ ends of CTG genomic repeats can induce large and uncontrolled sequence deletions, and the use of double guide-RNAs flanking the repeat expansion can lead to frequent sequence inversions, which remain toxic^20,21^. Since CRISPR-Cas9 approaches for manipulating DNA remain inefficient and controversial due to the risk of germline editing, an alternative approach targeting the repeat RNAs using deactivated Cas9 (dCas9) fused to an active ribonuclease has recently been shown to be effective in cells^22^. However, the efficiency of this approach for targeting nuclear CUG^exp^ RNA in DM1 patients could be limited by a number of factors, such as: 1) dCas9 retains affinity for DNA, which could compete for its binding to RNA, 2) the large size of dCas9 fusion proteins may limit their nuclear localization or delivery in vivo, 3) dCas9 utilizes short guide-RNAs which may increase the chance of off-target RNA cleavage and, 4) they require protospacer adjacent motifs (PAM) for efficient binding, which are not present in most human repeat expansion sequences. Thus, significant challenges remain for the development of efficient therapeutic strategies to target toxic nuclear RNAs in DM1 patients.

In contrast, bacterial-derived CRISPR-Cas13 systems bind specifically to RNA and function as endoribonucleases to cleave RNA, bypassing the risk of germline editing that is associated with DNA-binding CRISPR-Cas endonucleases23-25. Single residue mutations within the two nuclease domains of Cas13 generate a catalytically deactivated enzyme (dCas13), which retains programmable RNA binding affinity in mammalian systems without the requirement for PAM sequences for efficient targeting26. Due to their large size and lack of intrinsic localization signals, both Cas9 and Cas13 fusion proteins are inefficiently localized to the mammalian nucleus. In our recent pre-print manuscript describing the adaptation of CRISPR-Cas13 for inducing targeted RNA cleavage and polyadenylation (https://doi.org/10.1101/531111), we identified a non-classical nuclear localization signal (NLS) derived from the yeast Ty1 retrotransposon which promotes robust nuclear localization of Cas13. The powerful activity of the Ty1 NLS suggested that efficient targeting of nuclear RNAs could be achieved for either visualization or cleavage using CRISPR-Cas13.

To visualize nuclear RNAs using CRISPR-Cas13, we designed a fusion protein combining the catalytically dead Type VI-B Cas13b enzyme from *Prevotella sp*. *P5– 125* (dPspCas13b) with either a C-terminal enhanced Green Fluorescent Protein (eGFP) or red fluorescent protein, mCherry (Figure 1A). A 3x FLAG epitope tag (F) and Ty1 nuclear localization sequence (Ty1 NLS) were added to the N-terminus of the dPspCas13b fusion proteins to promote efficient nuclear localization, hereinafter referred to as *hilightR green* or *hilightR red* (Figure 1A). Toxic RNA foci are the cellular hallmark of DM1 and can be induced in many cell types by the expression of transgenes expressing expanded CUG repeats. To mimic the nuclear RNA foci found in patients with DM1, we utilized a vector containing 960 CUG repeats in the human DMPK 3’ UTR (DT960) (Figure 1B)^27^. The DT960 construct is sufficient to recapitulate RNA foci formation in cells and can be detected by Fluorescent In Situ Hybridization (FISH) using an antisense (CAG) repeat probe or with an mCherry-MBNL1 fusion protein (Supplementary Figures 1A and B). To target *hilightR* fusion proteins to CUG repeats, we designed a PspCas13b-compatible crRNA containing an antisense CAG repeat target sequence (CAGx9), which is predicted to hybridize with 9 CUG repeats (Figure 1C). Guided by the CAGx9 repeat crRNA, *hilightR green* and *red* were completely nuclear localized and highlighted nuclear RNA foci generated by the DT960 vector (Figure 1D and Supplementary Figure 1C). In contrast, co-expression of *hilightR* constructs with a non-targeting crRNA resulted in broad, un-localized nuclear fluorescence (Figure 1D and Supplementary Figure 1C).

**Figure 1.**
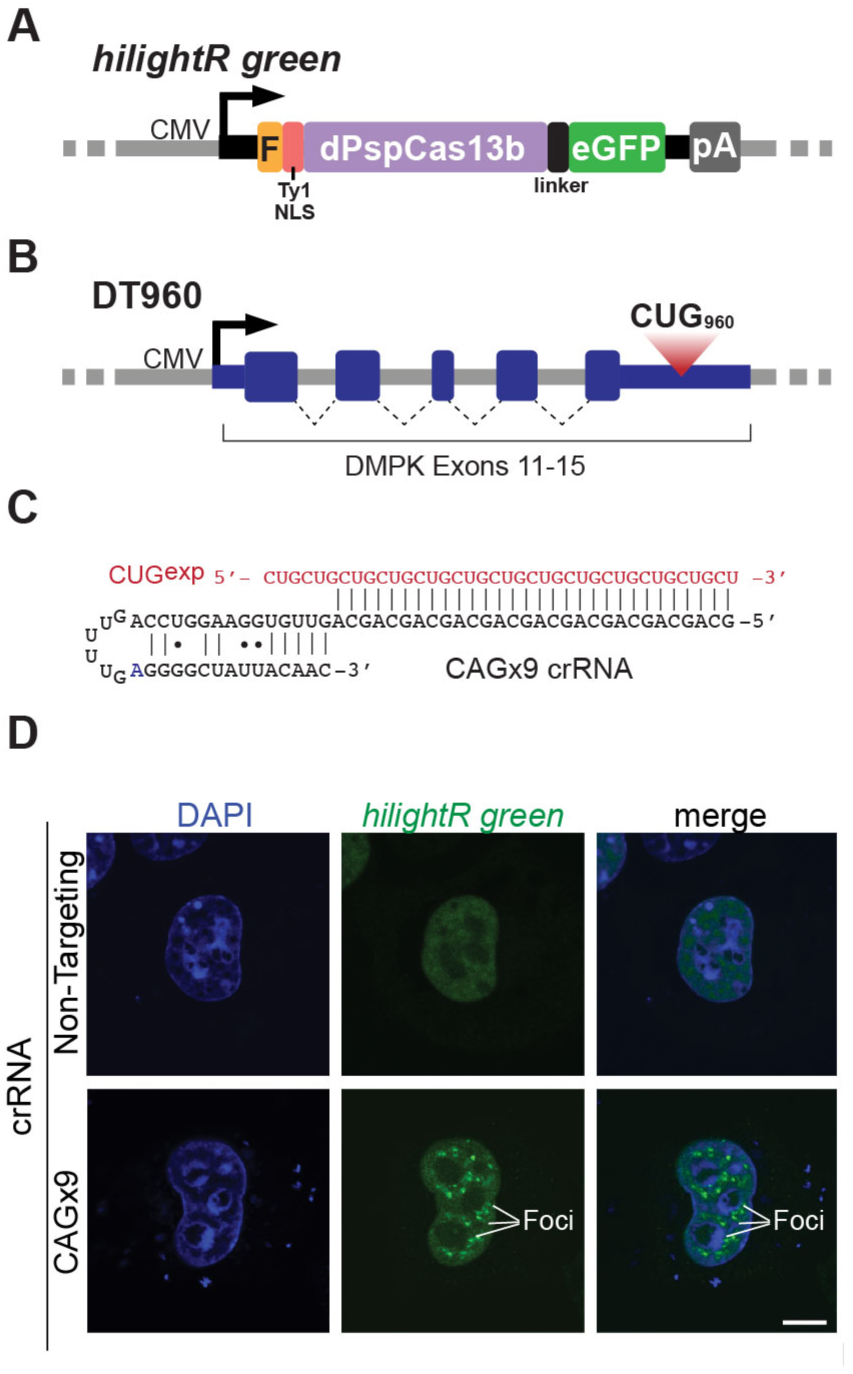
Development of robust nuclear localized CRISPR-Cas13 fusion proteins for the visualization of toxic RNA foci. (**A**) Design of catalytically dead PspCas13b (dPspCas13b) encoding an N-terminal 3x FLAG and Ty1 NLS and C-terminal eGFP. F – 3xFLAG epitope; NLS – Ty1 nuclear localization sequence; pA – SV40 polyadenylation sequence. (**B**) Diagram depicting the DT960 vector, which expresses a C-terminal genomic fragment of human DMPK (exons 11-15) encoding a 960 CTG repeat expansion in the 3’UTR. (**C**) Design of the CAGx9 crRNA and it’s predicted hybridization with CUG^exp^ RNA. (**D**) Representative images showing the cellular localization of *hilightR green* guided by either a non-targeting crRNA or the CAGx9 crRNA in COS7 cells expressing CUG^exp^ RNA. Scale bars, 10 µm.

Nuclear foci labeled with *hilightR green* co-localized with an Alexa Fluor 488-conjugated CAG oligonucleotide probe (AF488-CAGx7), detected using FISH (Figure 2A). Consistent with previous reports, nuclear foci labeled with *hilightR green* co-localized with MBNL1 protein, detected using an mCherry-MBNL1 fusion protein, and partially co-localized with splicing speckles, detected with an antibody specific for SC-35 (Figure 2B and Supplementary Figure 2). These results demonstrate that *hilightR* accurately targets CUG^exp^ RNA foci.

**Figure 2.**
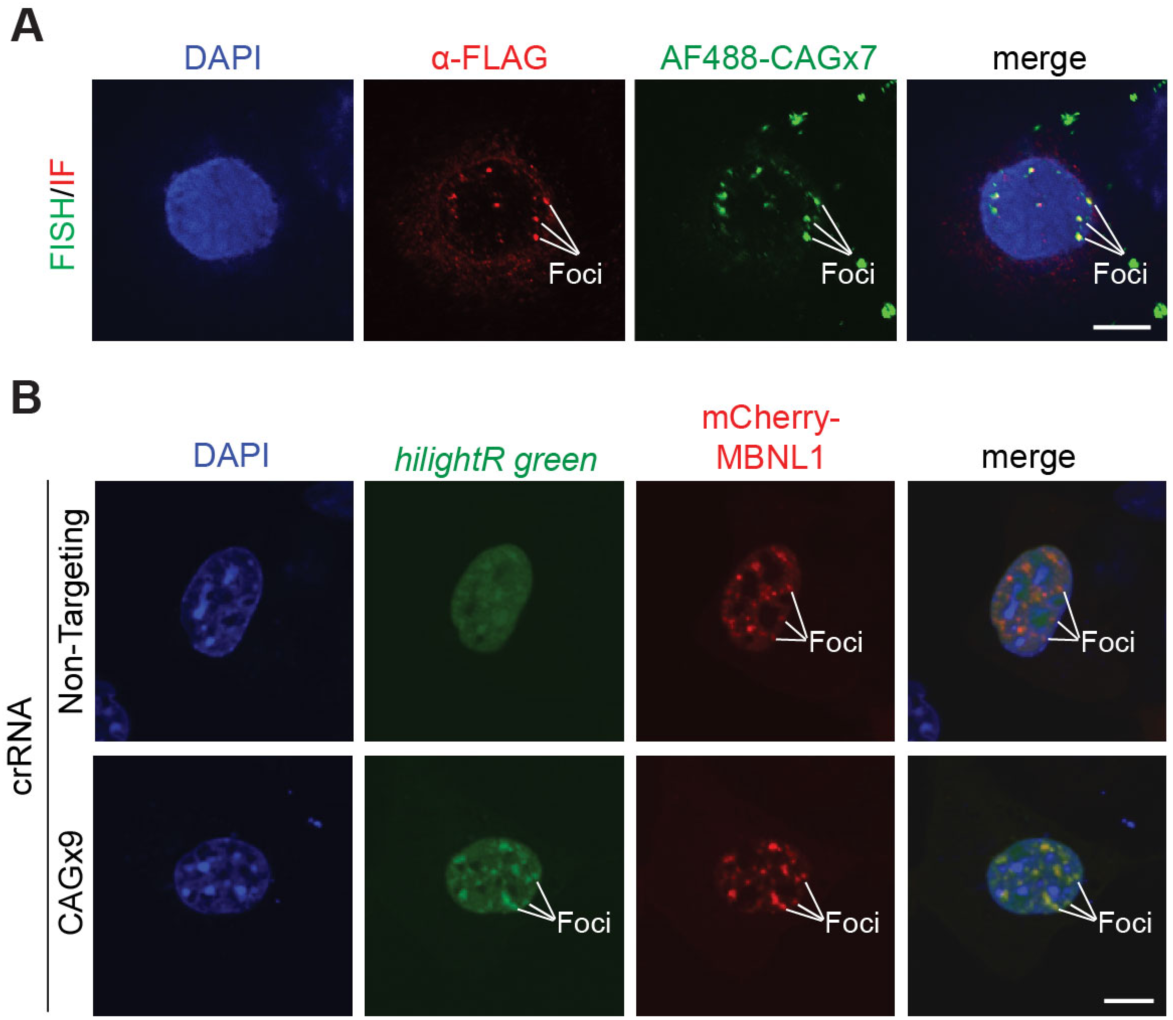
Co-localization of *hilightR green* with CUG^exp^ RNA foci and MBNL1. (**A**) Immunohistochemistry using an anti-FLAG antibody was used to detect *hilightR red*, targeted with the CAGx9 crRNA, which co-localized with CUG^exp^ RNA detected by FISH with an Alexa Fluor 488 CAGx7 probe. (**B**) *HilightR green* co-localized with mCherry-MBNL1 in COS7 cells expressing CUG^exp^ RNA foci when targeted with the CAGx9 crRNA, but not with a non-targeting crRNA. Scale bars, 10 µm.

The efficient nuclear targeting of CRISPR-Cas13 to CUG^exp^ RNA foci suggested it could be a useful for targeted cleavage of toxic CUG^exp^ RNA, using its inherent endoribonuclease activity. Cas13 has been shown to be useful for specific cleavage of mRNA transcripts in mammalian and plant cells^26,28,29^. To determine if Cas13 endoribonuclease activity is sufficient to cleave CUG^exp^ RNA foci, we modified the *hilightR green* fusion protein by reactivating PspCas13b’s catalytic mutations using site directed mutagenesis. Surprisingly, we found that activated *hilightR green* did not significantly reduce the number of RNA foci using the CAGx9 targeting crRNA, compared with a non-targeting guide-RNA (data not shown). However, activated PspCas13b containing the N-terminal Ty1 NLS, but lacking the C-terminal eGFP (herein referred to as *eraseR*), resulted in a significant reduction in the number and intensity of RNA foci, quantified using an mCherry-MBNL1 fusion protein (Figure 3A and B). Since target site flanking sequences can influence Cas13 nuclease activation, we tested CAGx9 crRNAs in two other reading frames (CAGx9-f2 and CAGx9-f3). *EraseR* guided by all three CAGx9 crRNAs resulted in significant reduction in the number and intensity of RNA foci per cell, compared to a non-targeting crRNA (Figure 3A and B). Additionally, catalytically dead PspCas13b containing an N-terminal Ty1 NLS and lacking the C-terminal eGFP did not significantly reduce the number and intensity of RNA foci, quantified using an mCherry MBNL1 fusion protein (Supplementary Figure 3). These date demonstrate for the first time that CRISPR-Cas13 is sufficient to degrade CUG^exp^ RNA foci.

**Figure 3.**
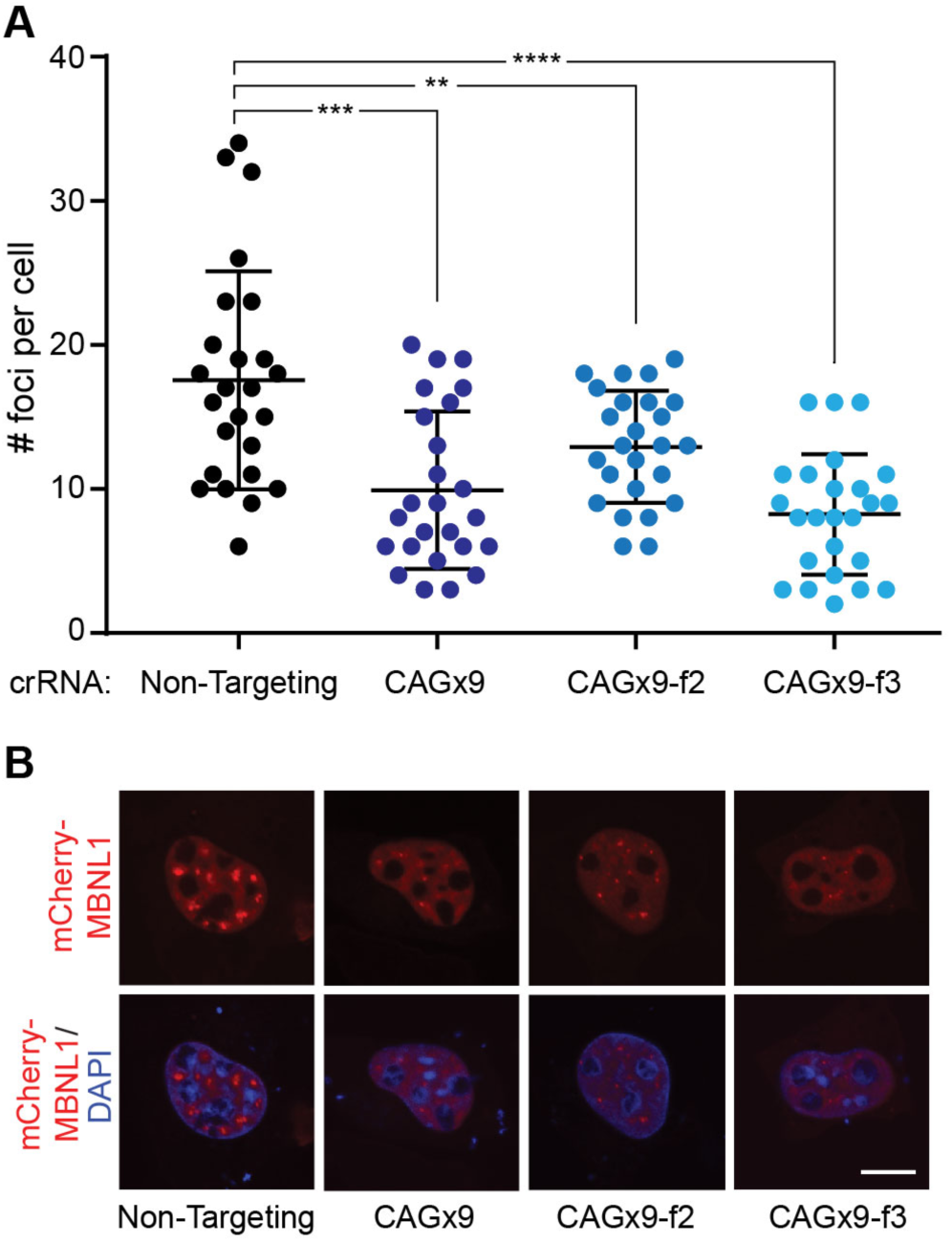
Degradation of toxic RNA foci by catalytically active PspCas13b. (**A**) Co-expression of active PspCas13b encoding a Ty1 NLS (*eraseR*) significantly decreased the number of RNA foci in cells expressing CUG^exp^ RNA, when targeted with all three CAGx9 crRNAs, detected using mCherry-MBNL1. (**B**) Representative micrographs of cells targeted by *eraseR* showing foci detected by mCherry-MBNL1, which are significantly decreased in number and appear fainter. Scale bars, 10 µm. ** = p-value < 0.01, *** = p-value < 0.001, **** = p-value < 0.0001.

While there are currently no available treatments for myotonic dystrophies or other human RNA repeat expansion disorders, the rapid development of CRISPR-Cas13 systems offer hope that targeted approaches to treat DM1 will soon be achievable. We have shown that Cas13b, localized by a powerful non-classical Ty1 NLS, can be used to efficiently target nuclear CUG^exp^ RNA foci for either visualization or targeted degradation in mammalian cells. The NLS used in these studies was derived from the yeast Ty1 LTR-retrotransposon (Ty1 NLS), which replicates its genome through a cytoplasmic intermediate^30^. As opposed to higher eukaryotes in which the nuclear envelope breaks down during cell division, the nuclear envelope in yeast remains intact during cell division and thus Ty1 requires active nuclear import for retrotransposition^31^. Interestingly, quiescent mammalian cells also retain a nuclear envelope, suggesting that the Ty1 NLS may be useful for targeting nuclear RNAs by Cas proteins in non-dividing cells.

The programmable nature of CRISPR-Cas13, through simple modification of crRNA target sequences, allow *hilightR* and *eraseR* to be easily adapted for the study and cleavage of other nuclear RNAs, or other repeat expansion disorders such as myotonic dystrophy type 2 (DM2), amyotrophic lateral sclerosis (ALS), huntington’s disease-like 2 (HDL2), spinocerebellar ataxias 8, 31 and 10 (SCA8, −31, −10) and fragile X-associated tremor ataxia syndrome (FXTAS). As with other approaches (ASOs or CRISPR-Cas) targeted to simple repeat sequences, there remains the potential for ‘coincidental’ cleavage of other human mRNA transcripts which contain these short repeat sequence motifs. Future studies will be necessary to evaluate any potential deleterious impacts resulting from coincidental cleavage events. Alternatively, directly targeting unique human DMPK transcript sequences for degradation by CRISPR-Cas13 or by other forms of RNA manipulation may offer additional approaches for the treatment of toxic RNA diseases.

## ACKNOWLEDGEMENTS

DT960 was a gift from Thomas Cooper (Addgene plasmid # 80412; http://n2t.net/addgene:80412; RRID: Addgene_80412).

## AUTHOR CONTRIBUTIONS

P.P., K.M.A. and D.M.A. designed the experiments. P.P., S.R.L, K.M.A., and D.M.A performed the experiments and analyzed data. P.P., K.M.A., and D.M.A generated the figures and wrote the manuscript.

## AUTHOR INFORMATION

The authors declare no competing financial interests. Correspondence and requests for materials should be addressed to D.M.A.

## METHODS

### Synthetic DNA and Cloning

The mammalian expression vector containing an N-terminal 3x FLAG and Ty1 NLS fused to dPspCas13b (described previously, https://doi.org/10.1101/531111) was modified to encode a C-terminal enhanced Green Fluorescent Protein (eGFP) or red fluorescent protein, mCherry. All crRNAs were designed to be 30 nucleotides in length and start with a 5’ G for efficient transcription from the hU6 promoter in pC00043^26^. The negative control non-targeting crRNA has been previous described^26^. To generate the mCherry-MBNL expression plasmid, the coding sequence of human MBNL1 was designed and synthesized for assembly as a gBlock (IDT, Integrated DNA Technologies) and cloned into the CS2mCherry mammalian expression plasmid^32^. Please see Supplementary Table 1 for cloned sequences. All plasmid insert sequences were verified by Sanger sequencing.

### Cell Culture, Transient Transfections and Immunohistochemistry

The COS7 cell line was maintained in DMEM supplemented with 10% Fetal Bovine Serum (FBS) with penicillin/streptomycin at 37°C in an atmosphere of 5% CO_2_. Briefly, 100,000 cells per well we seeded on glass coverslips in 6-well plates and transiently transfected after 24 hours using Fugene6 (Promega) according to manufacturer’s protocol. For *hilightR* experiments, cells were transfected with vectors encoding *highlightR*, crRNA, mCherry-MBNL1, and DT960 in a ratio of 2:2:2:1, and imaged twenty-four hours post-transfection. For *eraseR* experiments, cells were transfected with vectors encoding *eraseR*, crRNA, mCherry-MBNL1, and DT960 in a ratio of 2:2:2:1, and imaged forty-eight hours post-transfection. For immunohistochemistry, transiently transfected COS7 cells were fixed in 4% formaldehyde in DPBS for 15 minutes, blocked in 3% Bovine Serum Albumin (BSA) and incubated with primary antibodies in 1% BSA for 4 hours at room temperature. Primary antibodies used were anti-FLAG (Sigma, F1864) at 1:1000 and anti-SC-35 (Abcam, ab11826) at 1:1000. Cells were subsequently incubated with an Alexa Fluor 488 or 594 conjugated secondary antibody (Thermofisher) in 1% BSA for 30 minutes at room temperature. Coverslips were mounted using anti-fade fluorescent mounting medium containing DAPI (Vector Biolabs, H-1200) and imaged using confocal microscopy.

### Fluorescent In Situ Hybridization (FISH)

Twenty-four hours post-transfection, cells were fixed in ice cold 100% Methanol for 10 minutes at −20°C, then washed 2 times with DPBS and 1 time with Wash Buffer [2X SSC pH 7.0, 10% Formamide]. Cells were subsequently hybridized with probe in Hybridization Buffer [10% Dextran Sulfate, 2X SSC pH7.0, 10% Formamide] with a final probe concentration of 100 nM. Cells were hybridized overnight at 37°C. Cells were then washed one time in Wash Buffer at 37°C for 30 minutes, then mounted with VectaShield with DAPI (Vector Biolabs) on slides and imaged using confocal microscopy. The probe was a 21-mer DNA oligonucleotide (CAGCAGCAGCAGCAGCAGCAG) conjugated with a 5’ Alexa Fluor 488 dye and purified using HPLC (IDT, Integrated DNA Technologies).

## SUPPLEMENTARY DATA

## SUPPLEMENTARY FIGURES

**Supplementary Figure 1.**
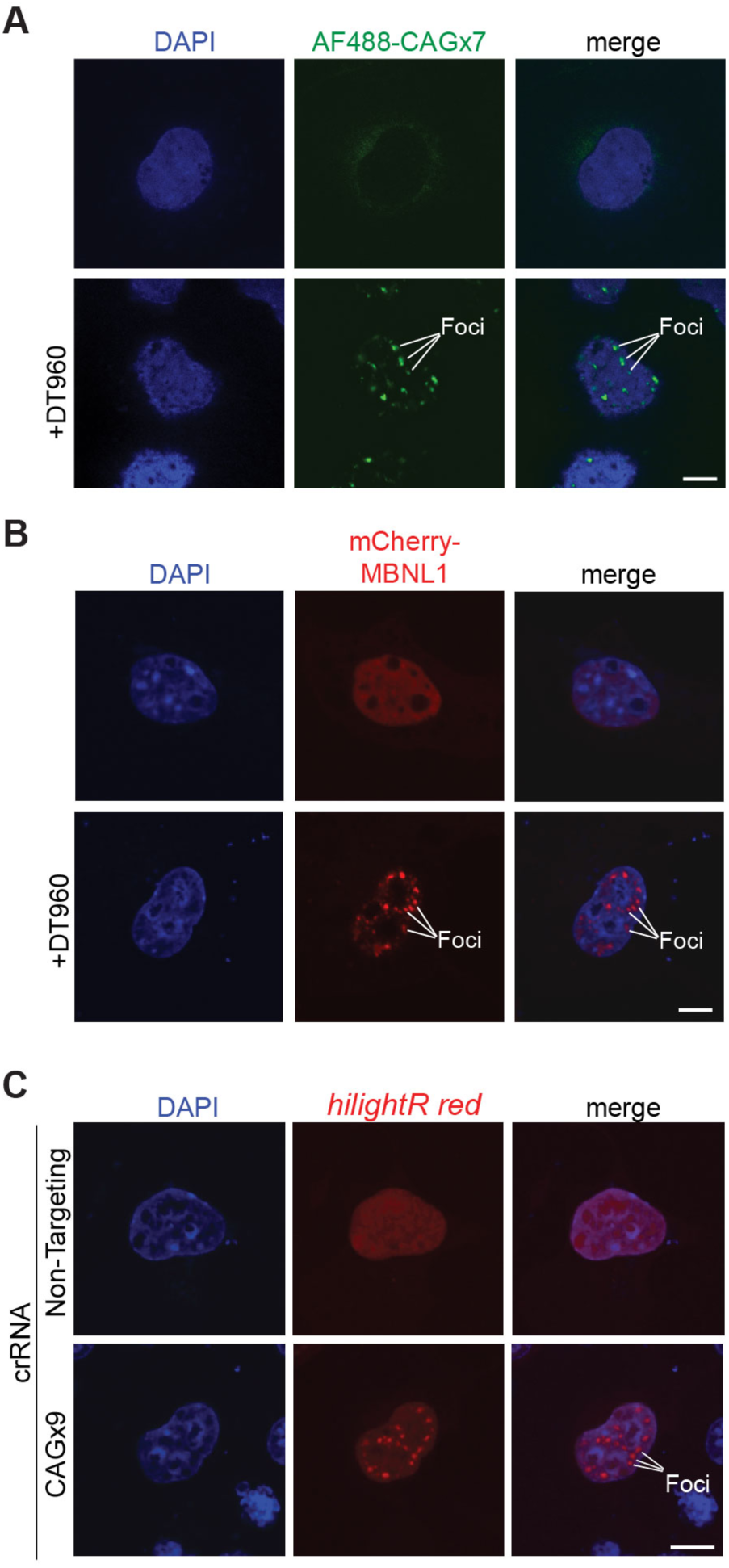
Detection of induced CUG^exp^ RNA foci in COS7 cells. (**A**) COS7 cells expressing 960 copies of CUG repeats induced RNA foci as detected using FISH with a CAG repeat antisense probe. AF488 – Alexa Fluor 488. (**B**) Expression of CUG^exp^ RNA induces the localization of MBNL1 to foci, as detected using an mCherry-MBNL1 fusion protein. (**C**) Localization of dPspCas13b-mCherry (*hilightR red*) guided by either a non-targeting or CAGx9 crRNA in COS7 cells expressing CUG^exp^ RNA. Scale bars, 10 µm.

**Supplementary Figure 2.**
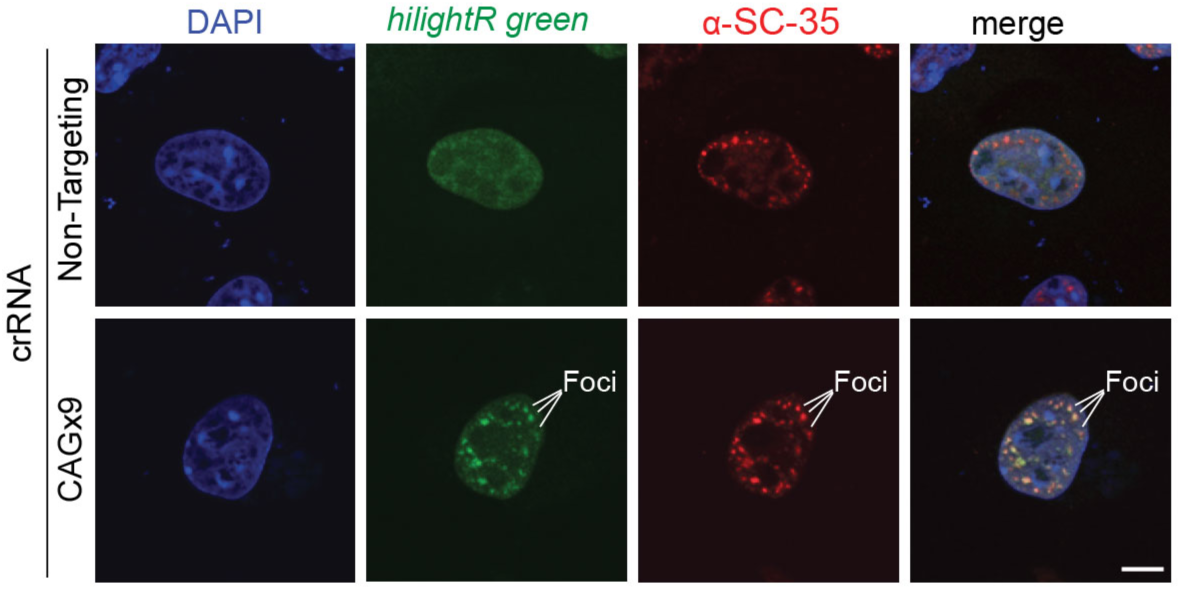
Co-localization of *hilightR green* with splicing speckles. In agreement with previous reports, CUG^exp^ RNA foci marked by *hilightR green* targeted with a CAGx9 crRNA, co-localized with splicing speckles, as detected using an anti-SC-35 antibody. Scale bars, 10 µm.

**Supplementary Figure 3.**
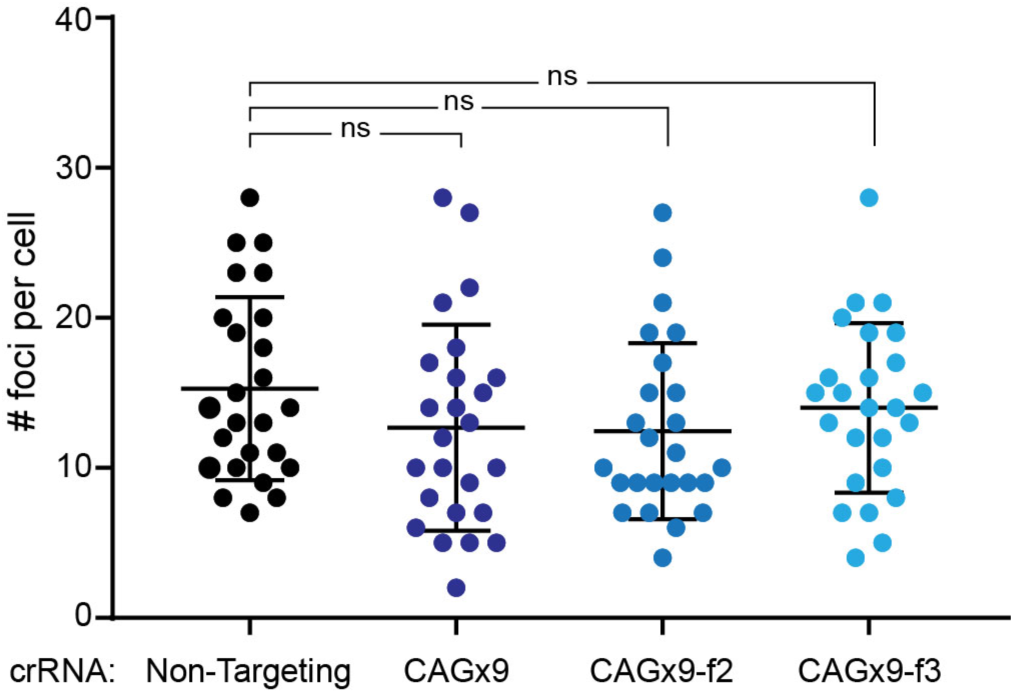
Catalytically dead Cas13 (dCas13) does not significantly reduce the number of CUG^exp^ RNA foci. Expression of dPspCas13b targeted with CAGx9 crRNAs does not significantly reduce the number of CUG^exp^ RNA foci per cell, as detected by mCherry-MBNL1. ns - not significant.

## SUPPLEMENTARY TABLE 1: Sequences

### PspCas13b crRNA target sequences (cloned into pC0043^26^)

*CAGx9*

5’-GCAGCAGCAGCAGCAGCAGCAGCAGCAGCA-3’

*CAGx9-f2*

5’-GAGCAGCAGCAGCAGCAGCAGCAGCAGCAGC-3’

*CAGx9-f3*

5’-GCAGCAGCAGCAGCAGCAGCAGCAGCAGCAG-3’

### QuikChange Site-Directed Mutagenesis primers

PspCas13b A133H

5’-ATGTACAGGGACCTGACCAACCACTACAAGACCTACGAGGAAAAG-3’

PspCas13b A1058H

5’-AAGATCCGGAACGCCTTCGATCACAACAATTACCCCGACAAAGG-3’

## hilightR green

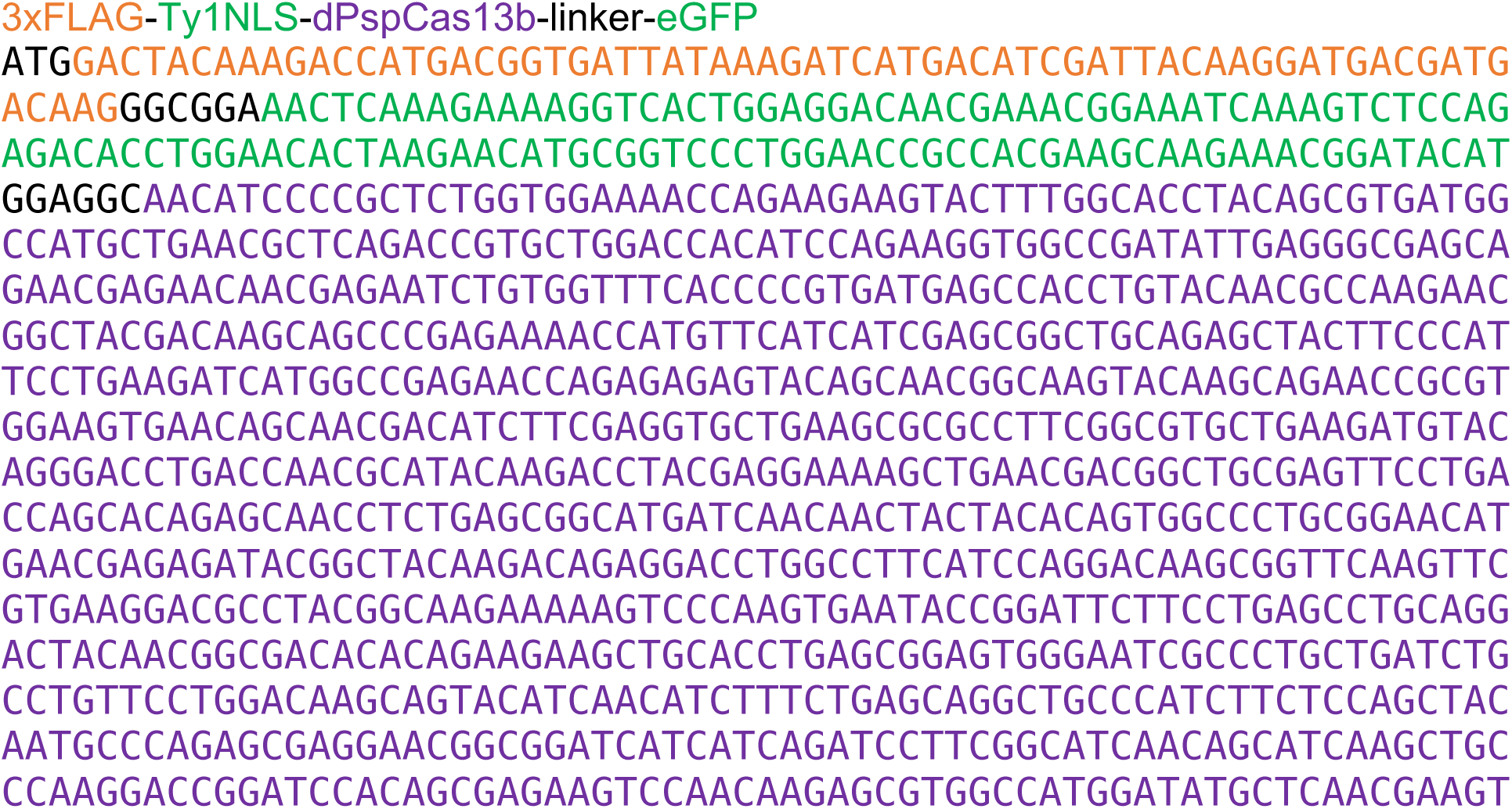
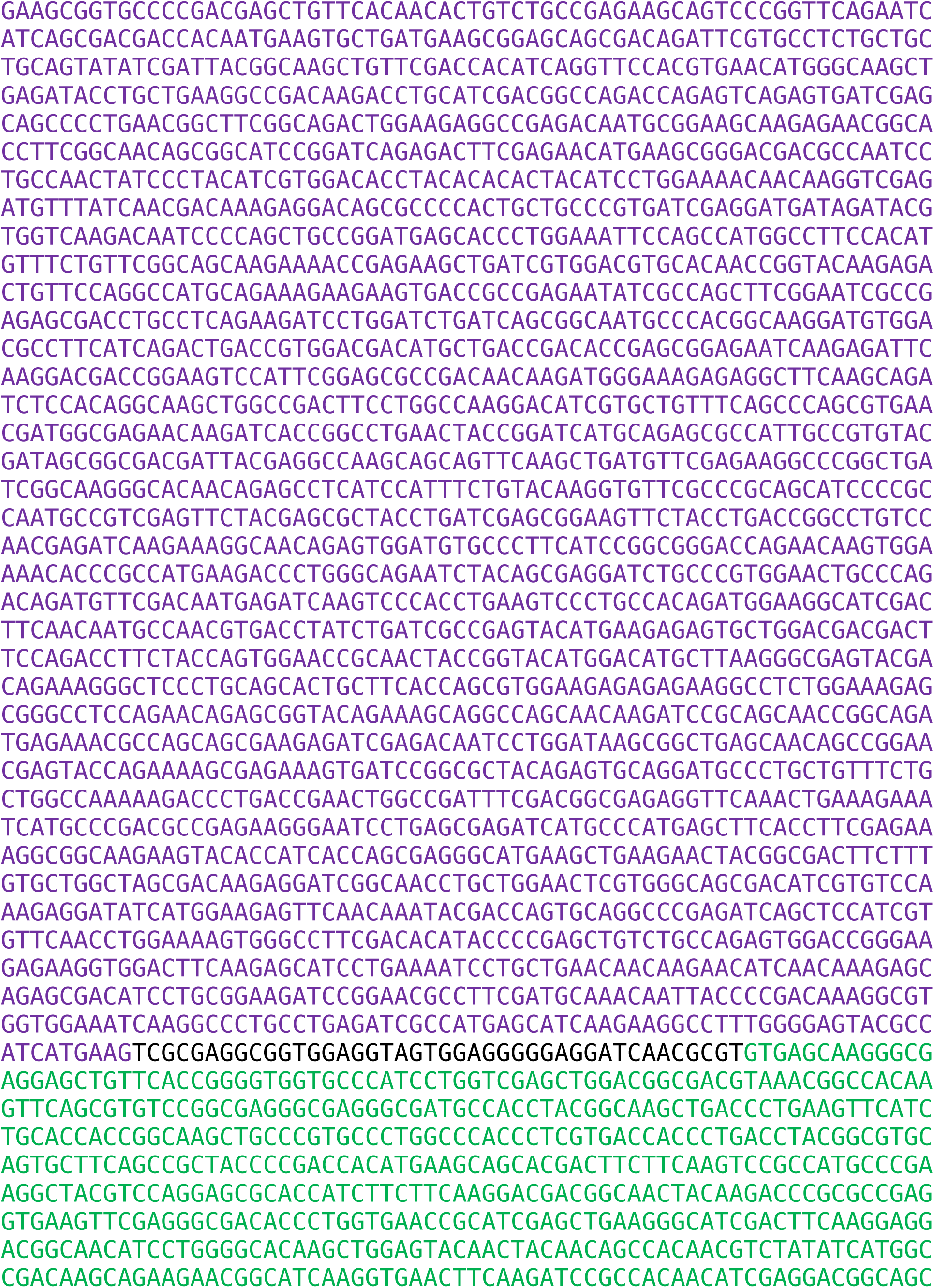
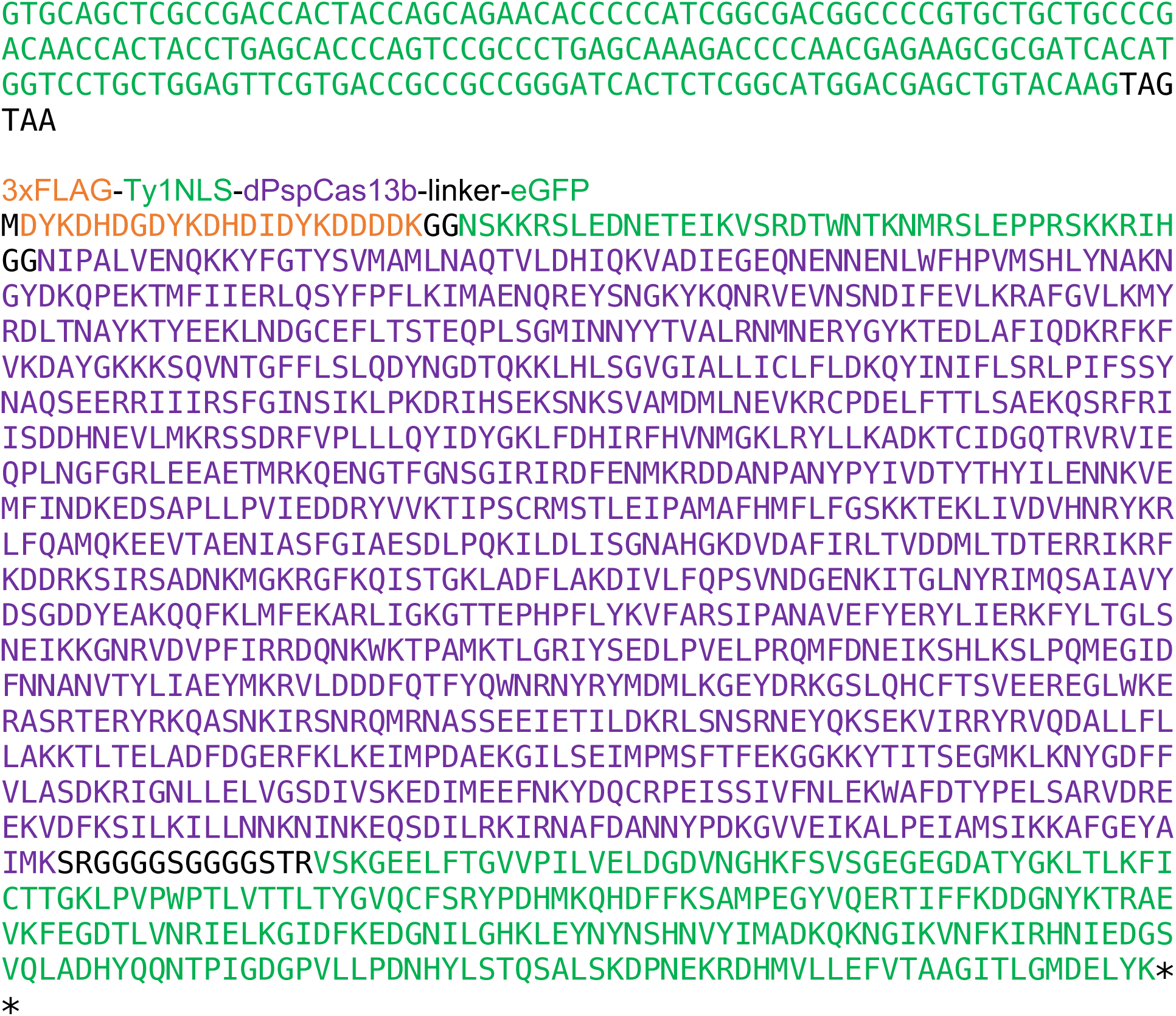

## hilightR red

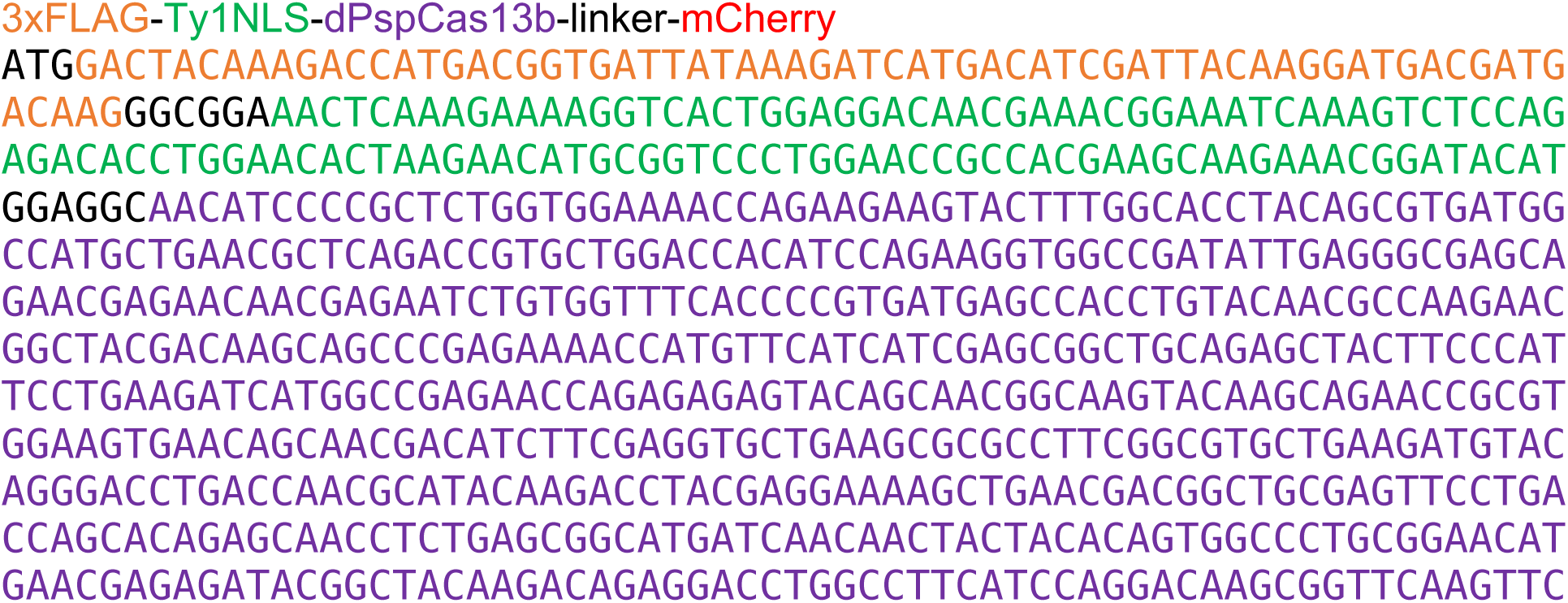
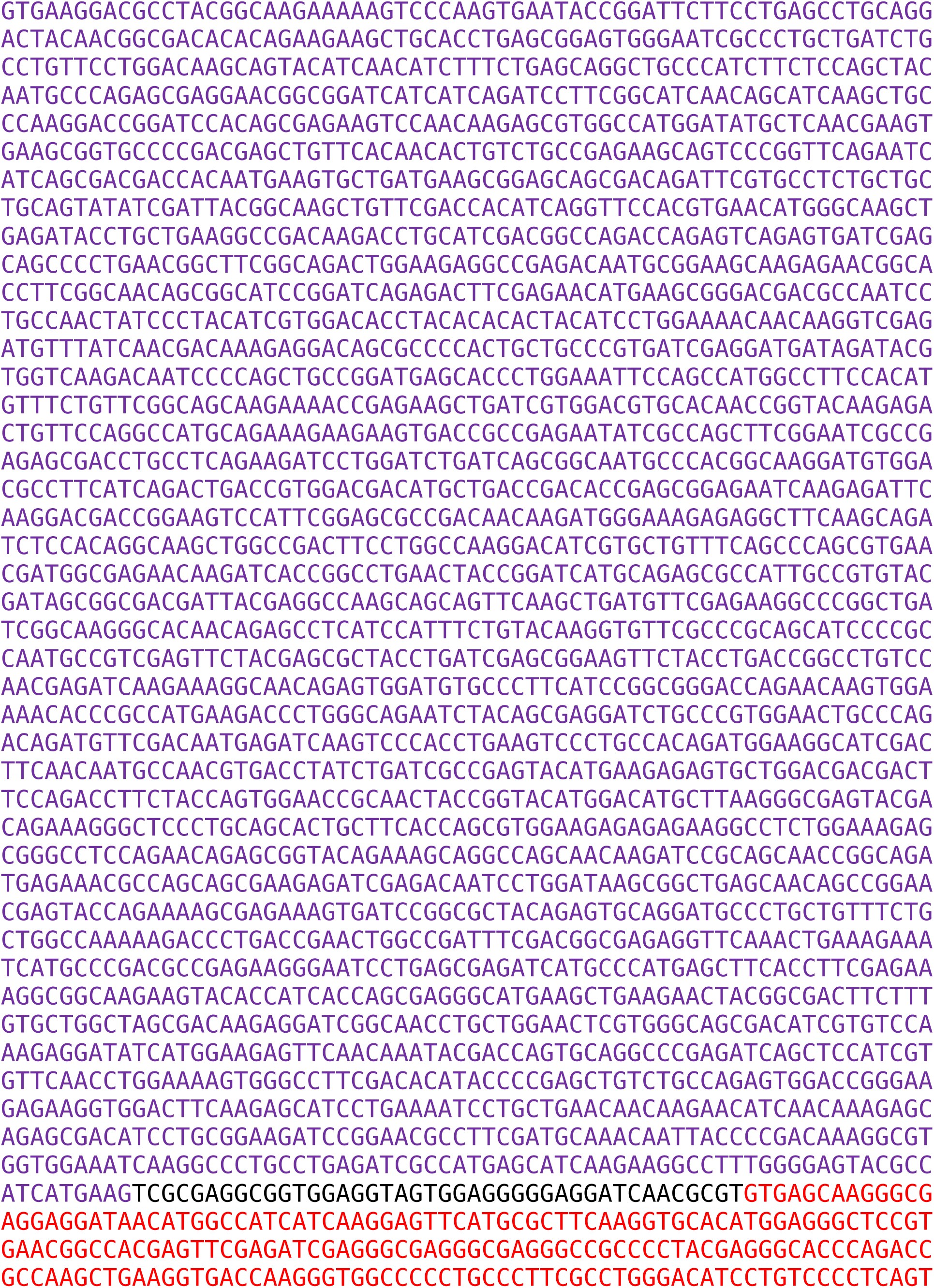
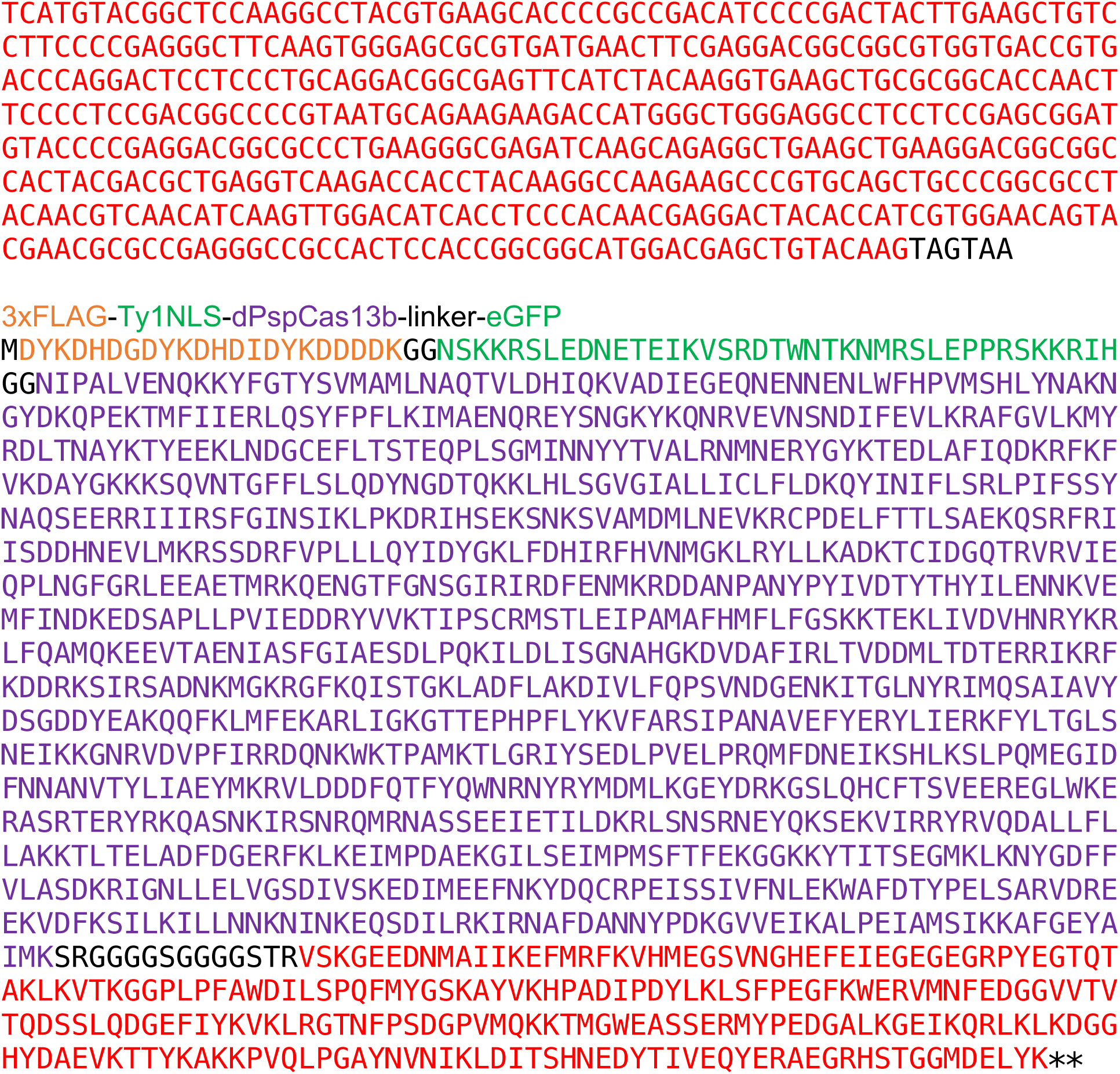

## CS2-mCherry-MBNL1

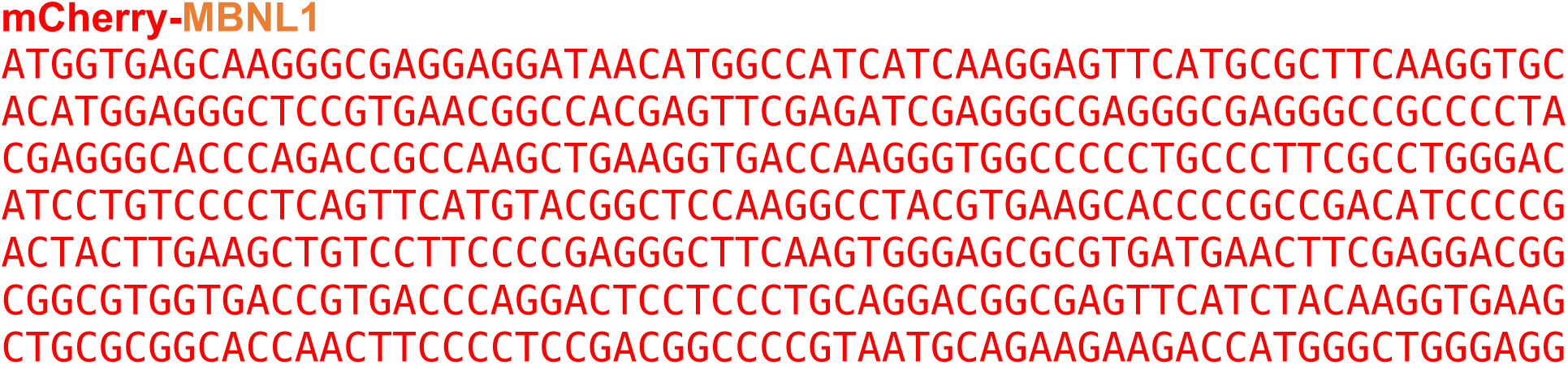
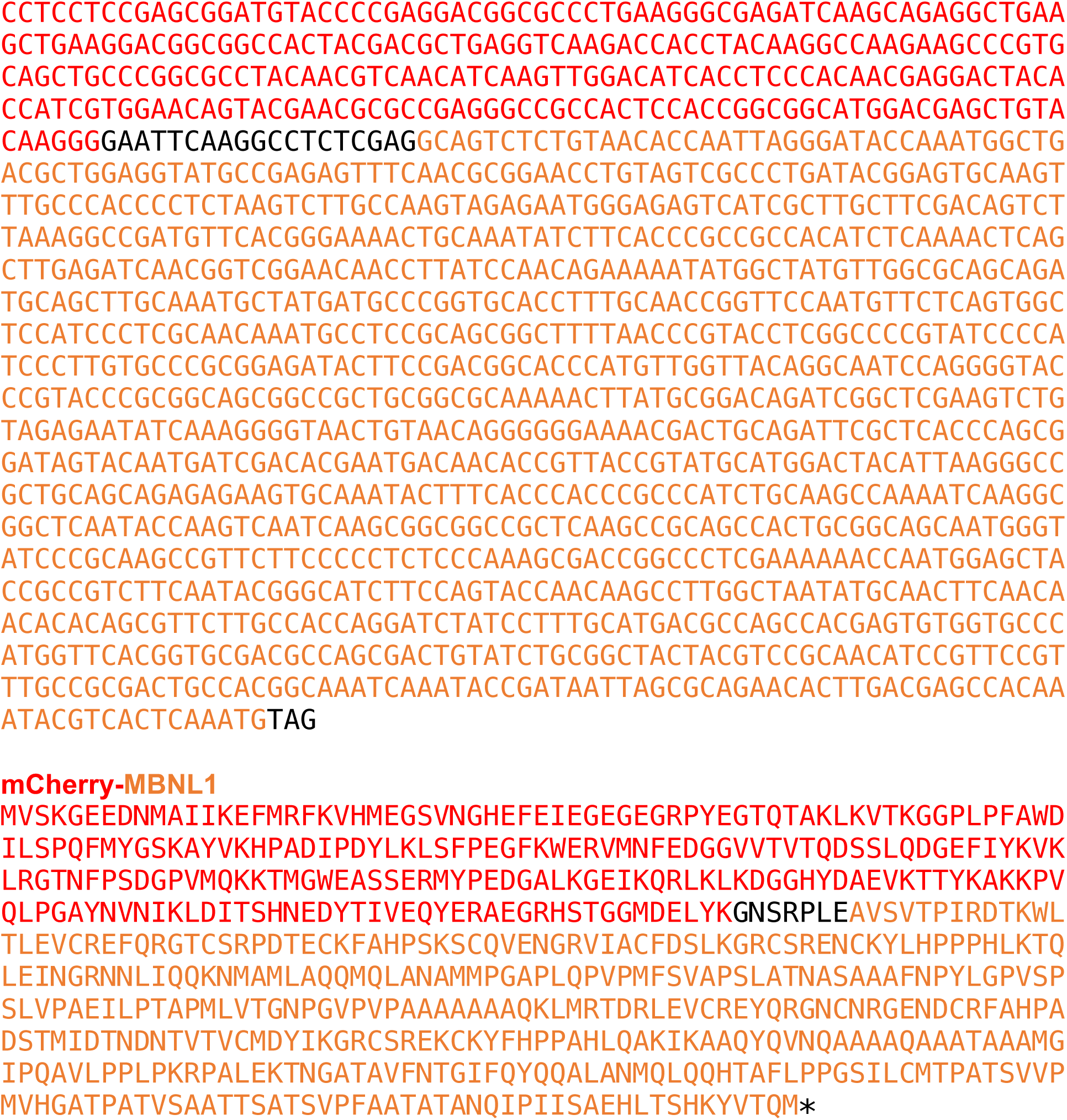

## eraseR

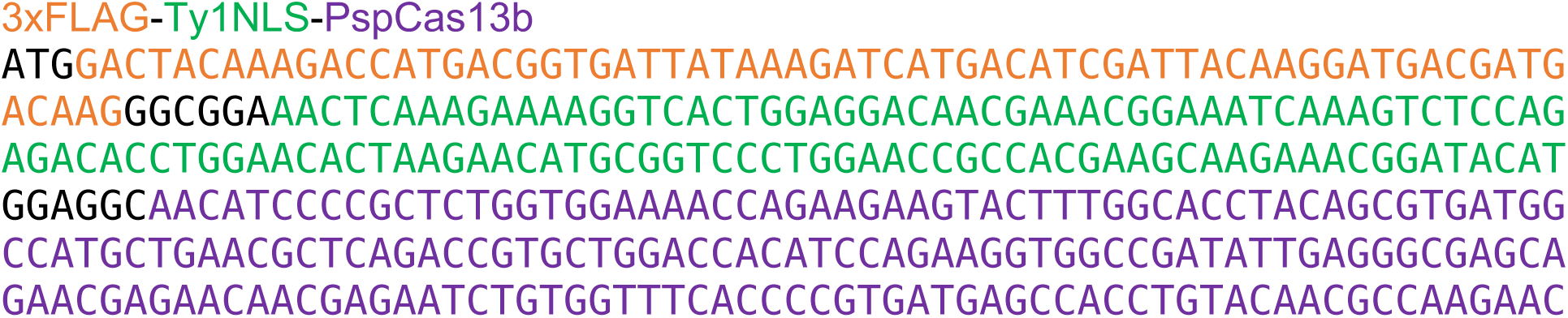
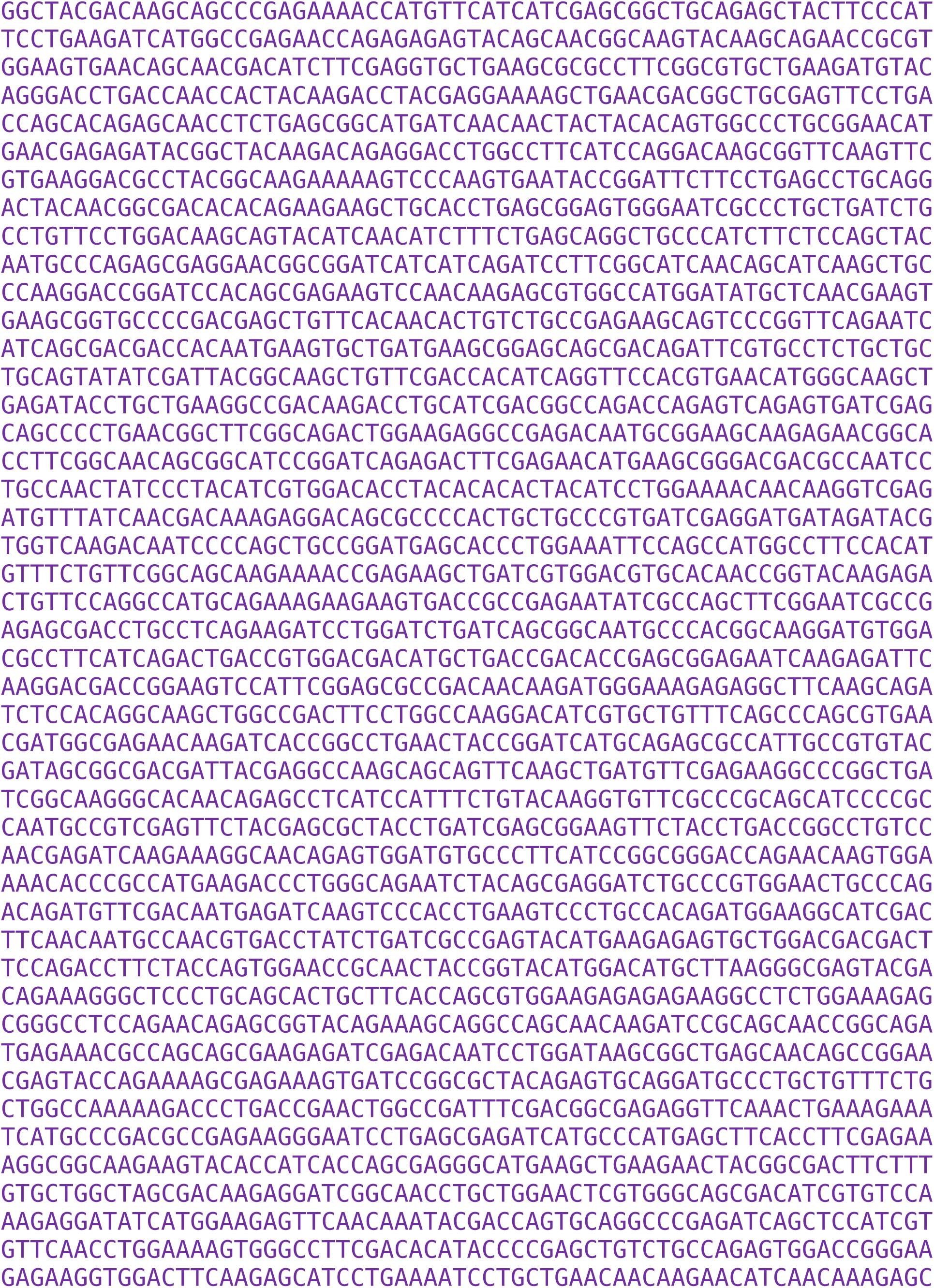
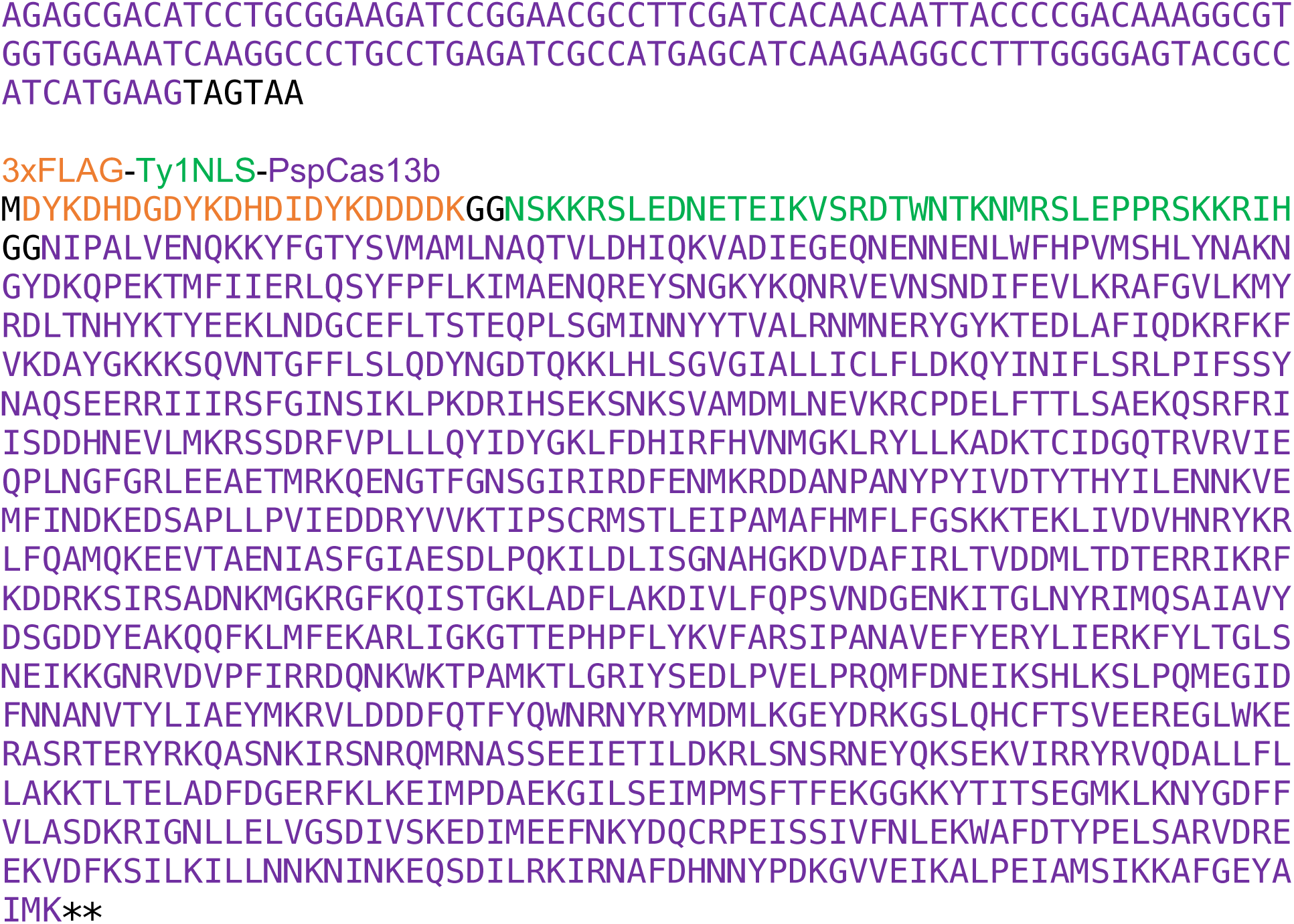

